# metapredict: a fast, accurate, and easy-to-use predictor of consensus disorder and structure

**DOI:** 10.1101/2021.05.30.446349

**Authors:** Ryan J. Emenecker, Daniel Griffith, Alex S. Holehouse

**Affiliations:** Department of Biochemistry and Molecular Biophysics, Washington University School of Medicine, St. Louis, MO, 63100, USA; Center for Science and Engineering Living Systems (CSELS), Washington University, St. Louis, MO 63130, USA; Center for Engineering Mechanobiology, Washington University, St. Louis, MO 63130, USA

## Abstract

Intrinsically disordered proteins and protein regions make up a substantial fraction of many proteomes where they play a wide variety of essential roles. A critical first step in understanding the role of disordered protein regions in biological function is to identify those disordered regions correctly. Computational methods for disorder prediction have emerged as a core set of tools to guide experiments, interpret results, and develop hypotheses. Given the multiple different predictors available, consensus scores have emerged as a popular approach to mitigate biases or limitations of any single method. Consensus scores integrate the outcome of multiple independent disorder predictors and provide a per-residue value that reflects the number of tools that predict a residue to be disordered. Although consensus scores help mitigate the inherent problems of using any single disorder predictor, they are computationally expensive to generate. They also necessitate the installation of multiple different software tools, which can be prohibitively difficult. To address this challenge, we developed a deep-learning-based predictor of consensus disorder scores. Our predictor, metapredict, utilizes a bidirectional recurrent neural network trained on the consensus disorder scores from 12 proteomes. By benchmarking metapredict using two orthogonal approaches, we found that metapredict is among the most accurate disorder predictors currently available. Metapredict is also remarkably fast, enabling proteome-scale disorder prediction in minutes. Importantly, metapredict is fully open source and is distributed as a Python package, a collection of command-line tools, and a web server, maximizing the potential practical utility of the predictor. We believe metapredict offers a convenient, accessible, accurate, and high-performance predictor for single-proteins and proteomes alike.

**Statement of Significance:** Intrinsically disordered regions are found across all kingdoms of life where they play a variety of essential roles. Being able to accurately and quickly identify disordered regions in proteins using just the amino acid sequence is critical for the appropriate design and interpretation of experiments. Despite this, performing large-scale disorder prediction on thousands of sequences is challenging using extant disorder predictors due to various difficulties including general installation and computational requirements. We have developed an accurate, high-performance and easy-to-use predictor of protein disorder and structure. Our predictor, metapredict, was designed for both proteome-scale analysis and individual sequence predictions alike. Metapredict is implemented as a collection of local tools and an online web server, and is appropriate for both seasoned computational biologists and novices alike.

## Introduction

While it is often convenient to consider proteins as nanoscopic molecular machines, such a description betrays many of their functionally critical features (1–3). As an extreme example, intrinsically disordered proteins and protein regions (collectively referred to as IDRs) do not adopt a fixed three-dimensional conformation (4–8). Rather, IDRs exist in an ensemble of different conformations that are in exchange with one another (9–11). Despite the absence of a well-defined structured state, IDRs are integral to many important biological processes (12, 13). As a result, there is a growing appreciation for the importance of disordered regions across the three kingdoms of life (6, 12, 14, 15).

A key first step in exploring the role of disorder in biological function is the identification of disordered regions. While IDRs can be formally identified by various biophysical methods (including nuclear magnetic resonance spectroscopy, circular dichroism, or single-molecule spectroscopy), these techniques can be challenging and are generally low throughput (16–18). As implied by the name, the “intrinsically” disordered nature of IDRs reflects the fact that these protein regions are unable to fold into a well-defined tertiary structure in isolation. This is in contrast to folded regions, which under appropriate solution conditions adopt macroscopically similar three-dimensional structures (19–21). The complexities of metastability in protein folding notwithstanding, this definition implies that this intrinsic ability to fold (or not fold) is encoded by the primary amino acid sequence (22–24). As such, it should be possible to delineate between folded and disordered regions based solely on amino acid sequence.

The prediction of protein disorder from amino acid sequence has received considerable attention for over twenty years, driven by pioneering early work by Dunker *et al*. (6–8, 25, 26). Since those original bioinformatics tools, a wide range of disorder predictors have emerged (27–30). Accurate disorder predictors offer an approach to guide experimental design, interpret data, and build testable hypotheses. As such, the application of disorder predictors to assess predicted protein structure has become a relatively standard type of analysis, although the specific predictor used varies depending on availability, simplicity, and scope of the question.

There are currently many disorder predictors that apply different approaches to predict protein disorder. These range from statistical approaches based on structural data from the protein data bank, to biophysical methods that consider local ‘foldability’, to machine learning-based algorithms trained on experimentally determined disordered sequences (31–38). However, using any individual predictor can be problematic; each predictor has specific biases and weaknesses in its capacity to accurately predict protein disorder, which can introduce systematic biases into large-scale disorder assessment (39). As such, an alternative strategy in which many different predictors are combined to offer a consensus disorder score has emerged as a popular alternative to relying on any specific predictor (40–44). Consensus scores report the fraction of independent disorder predictors that would predict a given residue as disordered - for example, a score of 0.5 reports that 50% of predictors predict that residue to be disordered.

While using consensus scores mitigates the limitations of any single predictor, calculating consensus scores is computationally expensive and necessitates installation of multiple distinct software packages. To alleviate this challenge, consensus disorder scores can be precomputed and held in online-accessible databases (42, 45–47). While precomputed scores are an invaluable resource to the scientific community their application is limited to a small subset of possible sequences. Furthermore, obtaining, managing, and analyzing large datasets of precomputed consensus predictions can be a daunting task, especially if only a subset of sequences are of interest.

To address these challenges we have developed a fast, accurate, and simple-to-use deep learning-based disorder predictor trained on pre-computed consensus scores from a range of organisms. Our resulting predictor, metapredict, is platform agnostic, simple to install, and usable as a Python module, a stand-alone command-line tool, or as a stand-alone web server. Metapredict accurately reproduces consensus disorder scores and is sufficiently fast such that for most bioinformatics pipelines, precomputation of disorder is no longer necessary, and disorder can be computed in real-time as analysis is performed. In addition to consensus disorder prediction, metapredict also provides structure confidence scores based on AlphaFold2-derived predictions of folding propensity, a related but complementary mode of sequence annotation. Metapredict can be installed in seconds, is incredibly lightweight, and has no specific hardware requirements. Taken together, metapredict is a high-performance and easy-to-use disorder predictor appropriate for computational novices to seasoned bioinformaticians alike.

## Materials and Methods

### Training metapredict using PARROT

To create metapredict, we used PARROT, a general-purpose deep learning toolkit developed for mapping between sequence annotations and sequence (48). PARROT was used to train a bidirectional recurrent neural network with long short-term memory (BRNN-LSTM) on the disorder consensus scores from the MobiDB database for each residue for all of the proteins in twelve proteomes (see supporting materials and methods for details) (**Figure 1**) (48–50). The eight disorder predictors used to generate the consensus scores in the MobiDB database were IUPred short (34), IUPred long (34), ESpiritz (DisProt, NMR, and X-ray) (31), DisEMBL 465 (28), DisEMBL hot loops (28), and GlobPlot (51). In total metapredict was trained using almost 300,000 individual protein sequences. For AlphaFold2-based predictions, the per-residue pLDDT score from nine different proteomes were used as input (see supporting materials and methods for details) (52, 53). The plDDT score reflects the confidence AlphaFold2 has in the local structure prediction.

**Figure 1.**
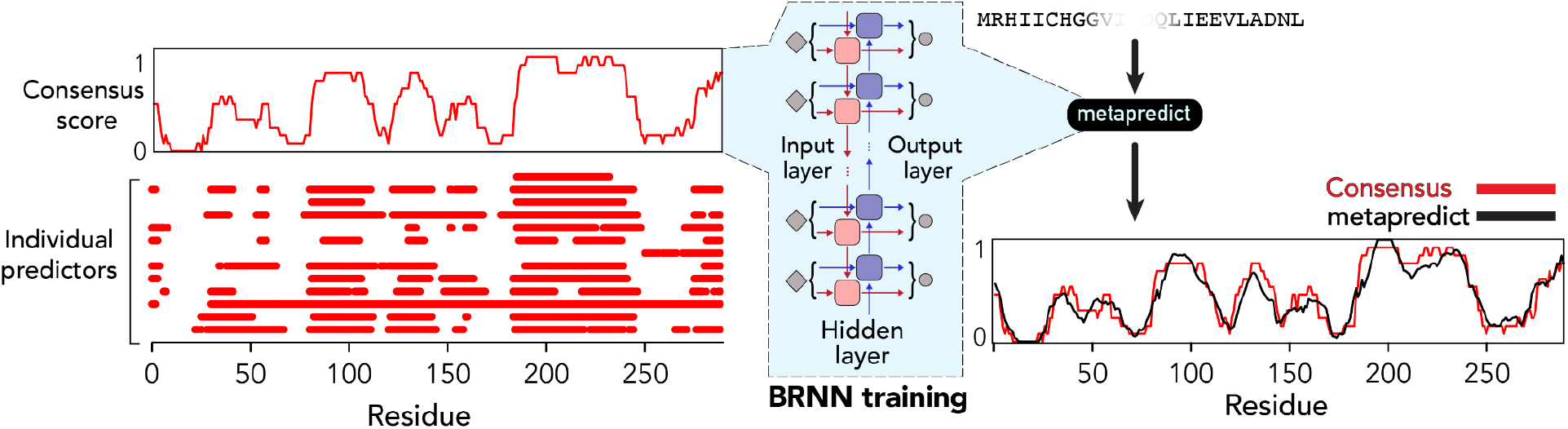
Overview of metapredict. Consensus scores are taken from 420,660 proteins distributed across 12 proteomes. Metapredict was developed by training a BRNN on this data, leading to a set of network weights that allow the prediction of any possible consensus sequence score.

Recurrent neural networks are well-suited for protein sequence machine learning tasks due to their ability to directly parse sequences of variable length without modification (54). Bi-directionality is a common modification of RNNs and is particularly relevant in the context of sequence-based prediction as it ensures that the entire local sequence (both N- and C-terminal) is accounted for when making the disorder prediction of a particular residue. Finally, LSTM networks are another common modification of RNNs that have seen widespread adoption in machine learning tasks because of their improved ability to retain long-range information over the course of training (50). Consequently bidirectional LSTMs have emerged as a powerful class of deep learning model for sequence-based predictions (48, 55–57).

To determine the optimal threshold to delineate disordered and ordered regions, we systematically varied the cutoff score used to classify IDRs (**Supplemental Figures 1-4**). This analysis revealed that a broad range of cutoffs (between 0.2 and 0.4) gave approximately equivalent performance, such that a cutoff of 0.3 offered a good balance between true positives and false negatives. As such, IDRs identified by metapredict with the default setting can be treated as relatively high-confidence, at the expense of missing some cryptic disordered regions.

### Usage and features

metapredict is offered in three distinct formats. As a downloadable package, it can be used either via a set of command-line tools or as a Python module. Command-line predictions include functionality to directly predict disorder from a UniProt accession, save disorder scores as a text file, and predict disorder for multiple sequences within a FASTA file. The Python module includes the ability to predict per-residue consensus disorder scores or delineate continuous IDRs. Complete documentation is available at http://metapredict.readthedocs.io/. In addition, we offer a web server appropriate for individual protein sequences, which is available at http://metapredict.net.

### Performance

On all hardware tested (which included a laptop from 2012), metapredict obtained prediction rates of ~7,000 to 12,000 residues per second (see supporting material for further details). A single 300-residue protein takes ~25 ms and the human proteome (20,396 sequences) takes approximately 21 minutes. Importantly and unlike some other predictors, the computational cost scales linearly with sequence length (**Supplemental figure 6**) (58).

## Results

### Evaluating metapredict accuracy in comparison to existing predictors

Given the large number of protein disorder predictors available, multiple groups have investigated different approaches to measure their accuracy (27, 59–61). Here, we used metrics from two recent studies, allowing us to compare directly with many previously evaluated predictors.

We first evaluated metapredict using the protocol developed for the Critical Assessment of protein Intrinsic Disorder experiment (CAID, 652 sequences). CAID is a biennial event in which a large set of protein disorder predictors are assessed using a standardized dataset and standardized metrics (27). CAID uses a curated dataset of 646 proteins from DisProt, a database of experimentally validated disordered regions (62). As such, evaluation using CAID’s standards offers a convenient route to benchmark metapredict against the state of the art.

In keeping with the assessments developed by CAID, we evaluated metapredict in its capacity to predict disorder across two distinct datasets (DisProt, DisProt-PDB) as well as its ability to identify fully disordered proteins (27). While DisProt contains only true positive disordered regions, DisProt-PDB contains true positive and true negative regions, making it more appropriate for robust validation of discriminatory predictors (27). To maintain consistency with CAID, we used the F1-score (defined as the maximum harmonic mean between precision and recall across all threshold values, Eq. 3) to compare metapredict against other predictors (27). The F1-score of metapredict in the analysis of the DisProt dataset ranked 12th highest out of the 38 predictors originally assessed (**Figure 2A**).

**Figure 2.**
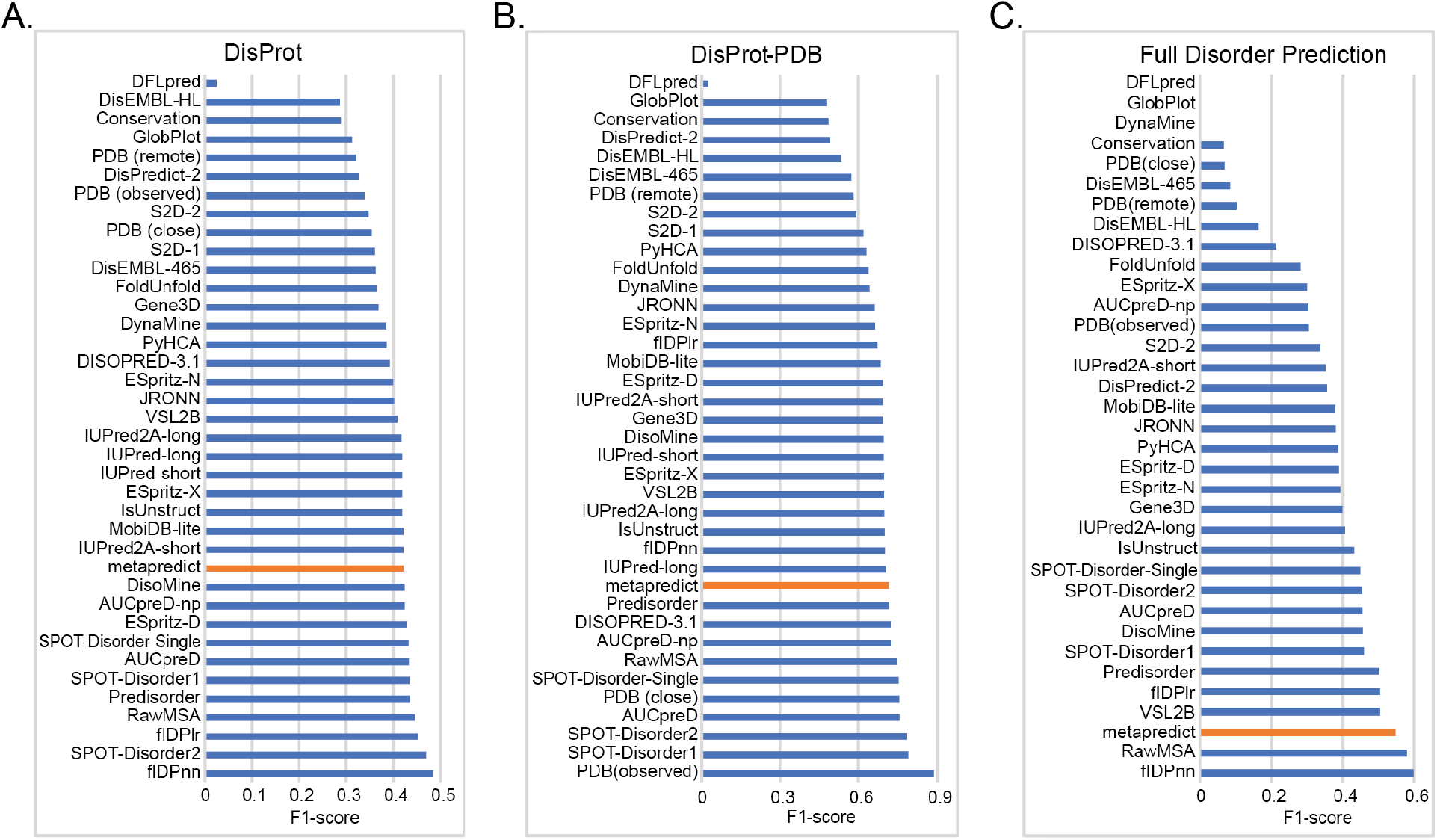
Evaluation of metapredict using CAID experiments. **(A)** F1-score for various predictors in examining their accuracy in predicting protein disorder from the DisProt dataset. **(B)** F1-scores for various predictors in examining their accuracy in predicting protein disorder from the DisProt-PDB dataset. **(C)** F1-scores for various predictors in predicting fully disordered proteins in the DisProt dataset. Values for all predictors in (A), (B), and (C) with the exception of those for metapredict (orange bar) were obtained from (27).

DisProt contains protein subregions that have been experimentally validated as disordered. However, as noted in the original study, it is possible, if not likely, that there are other subregions from those same proteins which, while not yet annotated as such, are in fact disordered (27). The DisProt-Protein Database (PDB) dataset addresses this limitation and includes only protein regions that are unambiguously annotated as either disordered or ordered, based on extant experimental data (27). In examining the performance of metapredict in predicting disorder on the DisProt-PDB dataset, we found that metapredict ranked 11th among all of the disorder predictors assessed (**Figure 2B**).

The last analysis that we carried out from the CAID experiment was the capacity of metapredict to identify fully disordered proteins. In this context, the CAID experiment considers something to be a fully disordered protein if the disorder predictor predicts 95% or more residues to be disordered (27). Metapredict ranked 3rd out of the disorder predictors examined in its capacity to identify fully disordered proteins (**Figure 2C**).

In addition to assessing metapredict via the CAID dataset, we also evaluated metapredict using the Chemical shift Z-score for assessing Order/Disorder (CheZOD), an alternative metric that provides a per-residue continuous value that experimentally quantifies disorder (see supporting information for more detail) (61). Similar to the CAID-based assessment, metapredict ranked on average 8th out of 23 predictors (**Supplementary Figure 1**).

While our assessment thus far is consistent with prior metrics, we worried that it lacked clear interpretability with respect to what these measures of accuracy mean for real protein sequences. To address this, we re-evaluated the CAID-derived predictions to compute an accuracy score that reflects the number of residues correctly predicted as folded or disordered per 100, using a Disprot-PDB-like dataset with any ambiguous residues excluded. **Fig. 3A** shows the resulting assessment and reveals that while the general order obtained from other methods is preserved (as expected), the difference between the best predictor and metapredict is on average two residues per 100.

**Figure 3.**
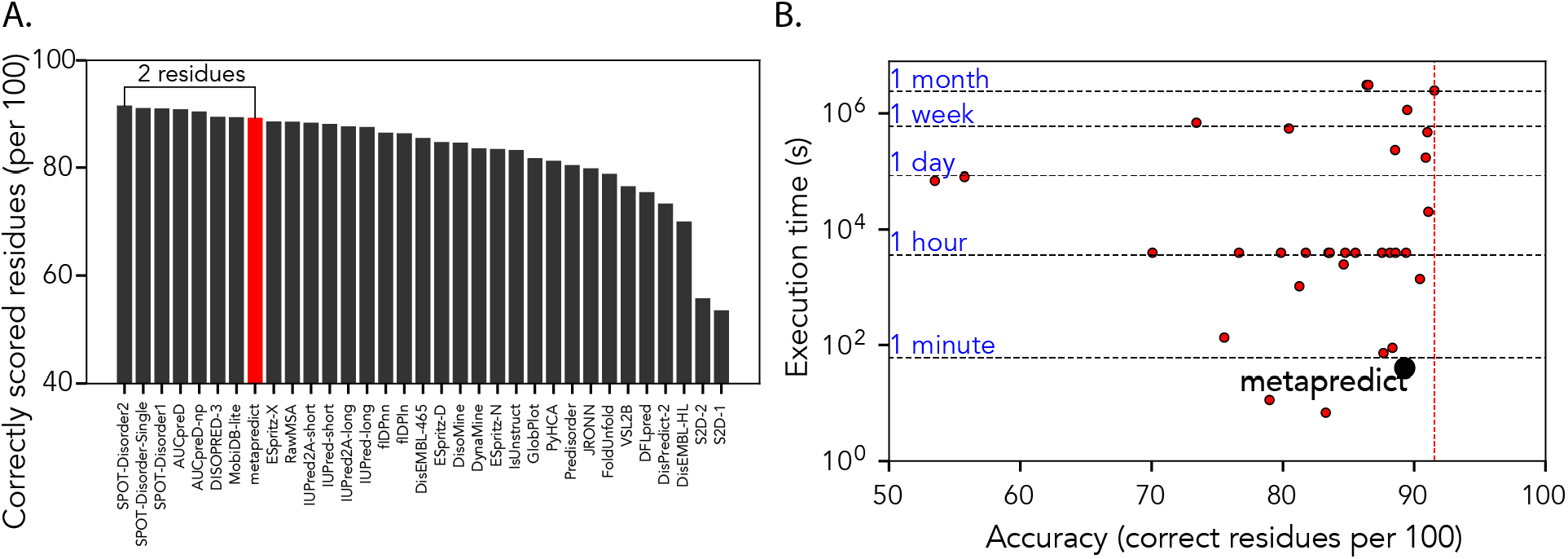
Accuracy and performance of metapredict. **(A)** Rank order of predictors in terms of number of correct residues per 100, assessed using true positive and true negative only (Disprot-PDB dataset). **(B)** Relative execution time for all predictors as evaluated in CAID over 652 independent sequences. Metapredict emerges as the third fastest predictor with a relative average loss in accuracy of 2 residues per 100 compared to the state-of-the-art (see all **Supplemental Figure S11**.)

### Evaluating metapredict execution time in comparison to existing predictors

Next, we considered how long metapredict takes to predict disorder compared to other predictors. AUCpreD was one of the top performing disorder predictors, and compared to several other top predictors was relatively easy to install. We evaluated the computational cost per-residue using the command-line version of metapredict. The time for AUCpreD-based disorder prediction scaled linearly with sequence length with approximately 0.3 seconds per residue (e.g. a 2151-residue protein takes ~14 minutes) (**Supplemental Fig. 2**). In contrast, no metapredict sequence took more than 0.9 seconds. In fact for single-sequence predictions, the main determinant of metapredict time was the time to load the trained network file (~0.6 seconds), which when predicting a FASTA file with multiple sequences is a fixed and negligible computational cost.

The CAID competition quantified execution times for 32 predictors using standardized hardware, providing a rigorous and complete assessment of relative performance. By scaling our hardware based on the CAID execution time scores for AUCPreD, we were able to compare accuracy and qualitative execution time of metapredict against all 32 predictors for the full CAID. While metapredict was ~2 residues per 100 less accurate than the top performing predictor, it took ~40 seconds to predict disorder for the full CAID dataset, compared to ~1 month. We tentatively suggest this difference in execution time compensates for the small drop in accuracy.

### Prediction of AlphaFold2 pLDDT prediction

In addition to direct disorder prediction and in response to the release of AlphaFold2-derived structure predictions for multiple proteomes, we developed a predictor for the per-residue confidence scores derived from the AlphaFold2 dataset (see supporting information for more detail) (52, 53). Formally, these scores reflect a predicted local difference test (pLDDT), such that metapredict offers a predicted prediction (i.e. a predicted pLDDT score) (**Fig. 4A**). Given the acquisition of structure can be considered the inverse of disorder, we expect (and observe) an anti-correlation between predicted structure confidence and disorder (**Fig. 4B, Supplemental Fig. S3**). We provide this feature as a complementary tool to aid in the interpretation of disorder scores, a feature that we anticipate will be useful when assessing ambiguous regions.

**Figure 4.**
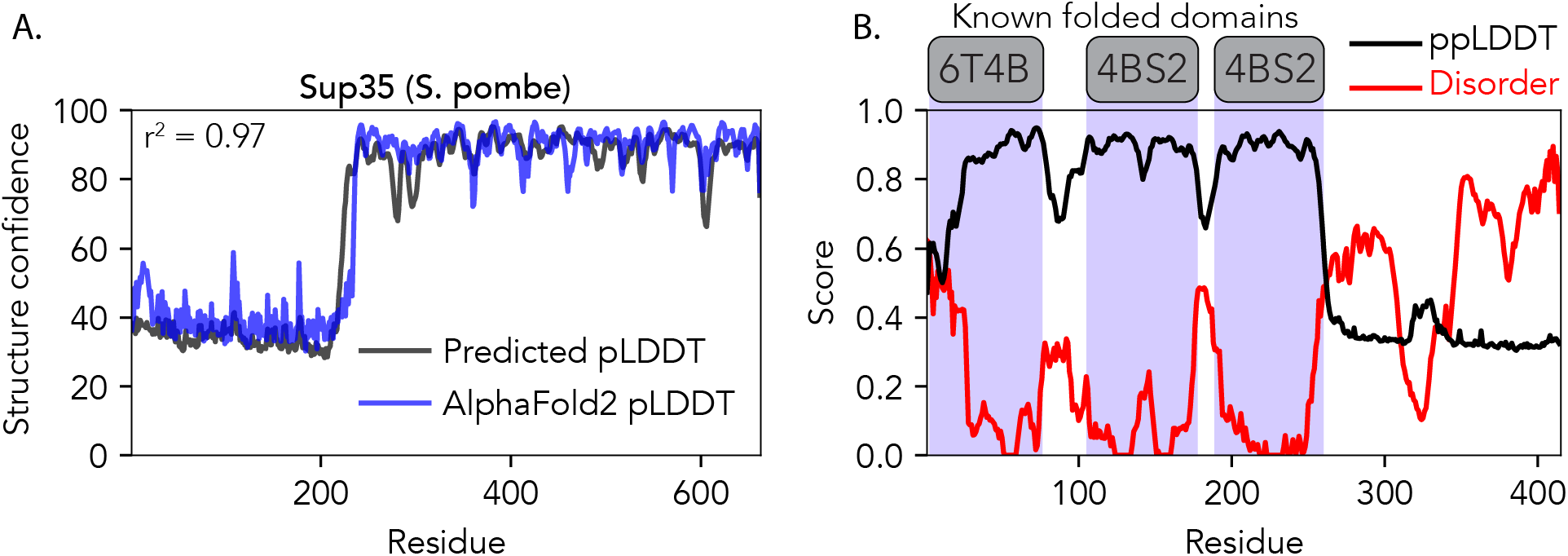
Metapredict also offers predicted structure confidence based on AlphaFold2. **(A)** Comparison of predicted pLDDT (blue) vs. actual pLDDT for the translational termination factor Sup35 from *Schizosaccharomyces pombe*. This sequence was not used in the training data, and is provided as a simple illustrative example of the agreement between the metapredict-derived prediction and actual AlphaFold2 pLDDT values **(B)**. Comparison of disorder (red), predicted pLDDT (ppLDDT, divided by 100 to place it on the same scale) and known folded domains (grey and blue) with associated PDB IDs shown for the human RNA binding protein TDP-43. The C-terminal disordered region is an experimentally verified IDR (63). Disorder and ppLDDT scores are anti-correlated, and correctly identify domain boundaries.

## Discussion

IDRs play many important roles in various biological processes (12, 13). An essential first step in the investigation of IDR function reflects the ability to identify IDRs within a protein sequence. Consensus disorder scores represent an attractive means by which to obtain high confidence disorder predictions that do not suffer from inaccuracies due to the limitations of any single disorder predictor. However, calculating disorder probabilities from many different predictors to generate a consensus score is cumbersome, technically challenging, and computationally expensive. To address this, we developed metapredict, a simple to use protein disorder predictor that accurately reproduces consensus disorder scores. While other consensus meta-predictors do exist, web-based access to these can be on the order of minutes-to-hours per sequence and where available, local access has operating-system dependencies making them poorly suited to cross-platform proteome-scale analysis (41, 64, 65). As such, we believe metapredict fills a niche that is currently unoccupied.

Metapredict makes use of a general approach in machine learning known as knowledge distillation. In knowledge distillation, a computationally cheap model is trained on data generated by one (or more) computationally expensive models, with a limited loss of accuracy (66, 67). This approach entirely detaches metapredict from either the computational cost or the computational complexity of other models, minimizing execution time, installation challenges, and limitations with respect to software or operating system dependencies.

In comparison to the other disorder predictors, metapredict tended to err on the side of false-negative predictions (where metapredict predicted something to be ordered when it was in fact disordered). As such, metapredict appears to possess a slight bias towards underestimating disorder, such that IDRs identified by metapredict can be considered reasonably high confidence. While metapredict is not the most accurate disorder predictor, we tentatively suggest the average error of 2 residues in 100 is relatively small. To aid in delineation between regions that may be ambiguous, the AlphaFold2 predicted structure confidence offers an orthogonal approach that provides additional discriminatory power.

### Features of metapredict

To further aid in the identification of *bona fide* contiguous disordered regions, metapredict contains a stand-alone function for extracting contiguous IDRs based on a threshold value applied to a smoothed disorder score and several additional parameters (**Supplemental Figures 4 - 7**). For this approach, we again found a threshold between 0.3 and 0.4 was optimal, and this method generally outperformed our prior more simple analyses. However, because other predictors did not use this approach for the classification of ordered or disordered regions, we chose to not use this function for our primary analyses in examining the accuracy of metapredict. Nonetheless, this suggests that metapredict can achieve even marginally higher accuracy in identifying IDRs and automates this procedure for the users, allowing boundaries between IDRs and folded domains to be automatically identified,greatly facilitating IDR-ome style analyses of datasets.

In addition to disorder prediction and in response to the recent release of AlphaFold2, metapredict offers an additional predictor of structure trained on AlphaFold2 data. The implications and application of AlphaFold2-derived predicted structure is an ongoing topic of investigation for many groups (68–71). While the absence of predicted structure cannot *necessarily* be taken to mean a region is disordered, there is a strong correlation and good reason to believe that for proteins in isolation, regions lacking high-confidence predicted structure may be disordered **(Supplemental Fig. 3)** (52, 53). As a final thought, predicting structure confidence using metapredict takes milliseconds, making this a potential screening tool for identifying high-confidence sequences of interest which could be investigate using the full AlphaFold2 methodology.

As a final note, an important feature in the distribution of software is ease of installation. Metapredict can be installed through a single terminal command (“pip install metapredict”), all dependencies are automatically included, and the metapredict package is just 3.8 MBs. This is in contrast to many other state-of-the-art predictors, which require large sets of additional tools (each of which must be separately installed) and hundreds of gigabytes of database files, and provide execution times on the order of minutes to hours per sequence. We believe metapredict offers an accurate, convenient and computationally efficient approach to *de novo* disorder prediction.

## Code and data availability

The code for metapredict can be found at: https://github.com/idptools/metapredict. Documentation is available at https://metapredict.readthedocs.io/. Fully processed sequences used for assessment (including sequences and scores) and code used for this manuscript are provided at https://github.com/holehouse-lab/supportingdata/. Metapredict can be installed directly from the Python Packaging Index using pip (i.e., “pip install metapredict”).

## Author Contributions

R. J. E. designed research, developed code, performed analysis, made figures, and wrote init manuscript. D.G. developed code, performed analysis, made figures, and wrote the manuscript. A.S.H. designed research, developed code, made figures, and wrote the manuscript.

## Acknowledgments

Funding for this project was provided by the Longer Life Foundation (an RGA/Washington University Collaboration) to A.S.H. and the William H. Danforth Plant Science Fellowship to R.J.E. We thank Ishan Taneja and Jeff Lotthammer for helpful comments on the manuscript. We thank Steven Boeynaems for the “motivation” to develop our web server. We thank DeepMind and EBI for providing all the AlphaFold2 data in such an accessible, robust, and timely manner. We also thank the entire Tosatto group, the ELIXIR Intrinsically Disordered Proteins Community, and HUPO-PSI Intrinsically Disordered Proteins Community (notably Silvio Tosatto, Zsuzsanna Dosztanyi, Damiano Piovesan, Wim Vranken and Norman Davey) for all the European-funded bioinformatics work that largely fuels the international IDP informatics space.

