## Supplemental Information for "metapredict: a fast, accurate, and easy-to-use predictor of consensus disorder and structure"

### Supplemental Materials and Methods

#### Evaluating metapredict using CheZOD scores

In line with previous work, we assessed how the continuous probability values of various predictors correlated with the CheZOD scores using the Pearson's correlation coefficient (Eq. 1) (Nielsen & Mulder, 2019). CheZOD scores increase with order and decrease with disorder. A Pearson's correlation coefficient of −1 would mean that a predictor is perfectly anti-correlated with the Z-scores, and 0 would mean that there is no correlation. As such, the results are displayed as the absolute value of the Pearson Correlation coefficient.

Based on the CheZOD, metapredict ranked 8th among the 23 predictors previously examined (**Supplemental Figure 1A**). We also calculated the area under a receiver operating characteristic curve (AUC). The receiver operating characteristic curve uses true positive values and false positive values to assess the accuracy of the various predictors, such that a perfect predictor would have an AUC of 1. Based on this assessment metapredict was ranked 11th out of the 23 predictors evaluated (**Supplemental Figure 1B**).

We next examined the accuracy of metapredict in predicting binary classification of either order or disorder. Previously, a CheZOD score of less than 8 was considered disordered (Nielsen & Mulder, 2019). When converting a metapredict score to binary classification, we considered any residue with a score of 0.3 or higher as disordered. For this analysis the Matthews Correlation Coefficient (MCC) was also calculated for each predictor (Eq. 2). The MCC uses a combination of false positives, false negatives, true positives, and true negatives in order to examine the accuracy of a classifier. We found that metapredict had the 8th highest MCC out of the predictors evaluated (**Supplementary table 1**).

#### Metapredict implementation and usage

metapredict is written in Python 3.7+ and uses PyTorch, with the initial network trained using PARROT (Griffith & Holehouse, 2021; Paszke et al., 2019).

We designed metapredict to be as flexible and user-friendly as possible. For example, metapredict can be used as a Python library (**Supplementary Figure 8A**), a stand-alone command-line tool (**Supplementary Figure 8B**) or a web server (http://metapredict.net) (**Supplementary Figure 8C**). Moreover, metapredict contains functionality to generate graphs or disorder scores from the command-line by directly inputting a single protein sequence, a UniProt accession number, or a FASTA file containing many sequences. Finally, in comparison to other predictors, which can take seconds, minutes or even hours per sequence, metapredict's computational performance makes it sufficiently fast that on-the-fly disorder prediction can be faster than reading pre-computed values from disk. It is this combination of accuracy, computational efficiency, ease of use, and flexibility that makes metapredict a convenient tool for any kind of disorder prediction, from single sequences to proteome-wide analyses.

To illustrate the ability of metapredict to predict consensus scores, **Supplementary Figure 9** shows the computed consensus scores and the analogous prediction for four proteins with IDRs. Across our datasets, we found that metapredict generally performed better than over two-thirds of the currently available disorder predictors examined, likely with a slight bias for false negatives when the default disorder threshold is applied.

For a list of features see the documentation at https://metapredict.readthedocs.io/.

**Evaluating metapredict disorder scores**

Evaluations and datasets for the CheZOD score analysis (116 sequences) can be found in (Nielsen & Mulder, 2019). Evaluations and datasets for the Critical Assessment of protein Intrinsic Disorder prediction (CAID) analysis (652 sequences) can be found in (Necci et al., 2021). For convenience, all sequences and scores used are also provided at https://github.com/holehouse-lab/supportingdata/. Details on results including additional statistical analyses and the raw performance scores for each predictor that was used for comparisons to metapredict can be found in the supporting materials and methods.

**Statistical analysis for evaluating the accuracies of disorder predictors**

Statistical analysis and predictor evaluation was carried out following the protocols described previously and reproduced here for completeness (Necci et al., 2021; Nielsen & Mulder, 2019).

Predictor evaluation is performed via the Pearson's Correlation coefficient ($R_p$), the Matthew's Correlation Coefficient (MCC), and the F1-score.

The Pearson's Correlation Coefficient ($R_p$) is calculated as,

$$R_p = \frac{\sum(x_i - x_a)(y_i - y_a)}{\sqrt{\sum(x_i - x_a)^2 \sum(y_i - y_a)^2}} \quad \text{(Eq. 1)}$$

where $x_i$ is the value of the current predicted disorder value for a residue in a sequence, $x_a$ is the mean predicted disorder value for the residues in a sequence, $y_i$ is the actual disorder value (specifically the CheZOD score) of the current residue, and $y_a$ is the mean actual disorder value.

The Matthew's Correlation Coefficient (MCC) is calculated as,

$$MCC = \frac{(TP \times TN) - (FP \times FN)}{\sqrt{(TP + FP) \times (TN + FP) \times (TP + FN) \times (TN + FN)}} \quad \text{(Eq. 2)}$$

Where true positives (TP) are the number of times a disorder predictor predicts a disordered residue to be disordered, true negatives (TN) are the number of times a predictor does not predict something to be disordered when it is not disordered, false positives (FP) are the number of times a predictor predicts a residue to be disordered when it is in fact not disordered, and false negatives (FN) are the number of times a predictor predicts a residue to not be disordered when it is in fact disordered.

Finally, the F1-score is calculated as

$$\text{F1-score} = \frac{TP}{TP + (0.5 \times (FP + FN))} \quad \text{(Eq. 3)}$$

Where TP, FP, and FN are defined above.

**Metapredict training of disorder predictor**

The metapredict disorder prediction was trained using PARROT with slight modifications to the default settings (Griffith & Holehouse, 2021). We set the number of training epochs, which defines the number of times the complete dataset is assessed and used to update the network parameters, to 100. Increasing the number of epochs further gave a negligible improvement in performance. The PARROT data type was set to 'residues' and the number of classes was set to '1' (for regression). The learning rate (--learning-rate), which is a parameter that alters the rate that the model updates weights after each round of back-propagation, was set to 0.001. The number of layers (--num-layers), which is the number of layers in the network between the input layer and the output layer, was set to 1. The hidden vector size (--hidden-size), which is the size of hidden vectors within the BRNN, was set to 5. The batch size (--batch), which is the number of sequences processed at the same time, was set to 32. For training, validation, and testing, 70% of the data was used for training, 15% of the data was used for validation, and 15% of the data was used for testing.

The proteomes for which consensus disorder scores were available at the time of training were: *Danio rerio* (UP000000437, 43,841 proteins), *Gallus gallus* (UP000000539, 25,238), *Mus* *musculus* (UP000000589, 44,470 proteins), *Drosophila melanogaster* (UP000000803, 21,114 proteins), *Dictyostelium discoideum* (UP000002195, 12,733 proteins), *Canis lupus familiaris* (UP000002254, 45,089 proteins), *Saccharomyces cerevisiae* (UP000002311, 6,049 proteins), *Rattus norvegicus* (UP000002494, 29,090 proteins), *Homo sapiens* (UP000005640, 66,835), *Arabidopsis thaliana* (UP000006548, 39,342 proteins), *Sus scrofa* (UP000008227, 49,792 proteins), and *Bos taurus* (UP000009136, 37,367 proteins). These numbers reflect protein sequences composed of the 20 standard amino acids only. Cross-referencing the training dataset (70% of the total sequences) taken from these proteomes against the assessment databases used (CheZOD and CAID) identified 28/116 from CheZOD and 451/652 from CAID databases in a total training set of ~295,000 sequences. These proteomes and the associated consensus disorder scores were obtained from MobiDB, but were originally curated by UniProt (Acids Research & 2021, 2021; Piovesan et al., 2021; UniProt Consortium, 2019).

**Metapredict performance**

Metapredict has no specific hardware requirements and performs well across all set ups tested. Hardware tested included an Unbuntu-running Dell desktop (Ubuntu 18.04 with Intel(R) Core(TM) i9-9900 CPU @ 3.10GHz CPU, 32 GBs DDR4 RAM, with a Toshiba 512 GB SSD (KBG40ZNS512G), a 2020 Apple Mac Mini (16 GB unified memory, Apple M1 processor), a 2019 16-inch MacBook Pro (64 GB 2667 MHz DDR4 RAM, 2.3 GHz Intel Core i9 processor, Intel UHD Graphics 630 integrated graphics), and a 2012 MacBook Pro (2.9 GHz Intel Core i7 processor, 8 GB 1600 MHz DDR3 RAM, Intel HD Graphics 4000 1536 MB integrated graphics). Importantly, as evident by our testing on a 2012 MacBook Pro (which scored at 7,238 residues per second), metapredict does not require a high-end modern computer to be fast. Even on the most basic virtual machine available, (Ubuntu 18.04 with single virtual Intel CPU (2.4 GHz), 1 GB DIMM memory, 20 GB SSD), metapredict performs at ~6000 residues per second.

To compare AUCPreD vs. metapredict (**Supplementary Figure S1**) predictions were performed on a desktop machine running Ubuntu 18.04 with an Intel(R) Core(TM) i9-9900 CPU @ 3.10GHz CPU, 32 GBs DDR4 RAM, with a Toshiba 512 GB SSD (KBG40ZNS512G). To ensure a fair comparison, each sequence was isolated and placed in its own FASTA file, and the predictor ran on each file independently. In reality, if one had multiple sequences, placing them into a single FASTA file would be a more efficient approach to minimize the amount of time reading from the filesystem. While the effective contribution of file read time is negligible for AUCPreD, in all cases for metapredict it is the dominant determinant of execution time.

To compare metapredict execution time against other predictors based on times reported in the CAID experiment, we used per-sequence execution times for AUCPreD on our local hardware and on the CAID hardware to calibrate and approximate conversion factor. A more detailed explanation of this process is described in https://github.com/holehouse-lab/supportingdata/tree/master/2021/emenecker_metapredict_2021/performance/metapredict_on_caid_disprot

**AlphaFold2 pLDDT predictor data**

We obtained pLDDT scores for every AlphaFold2 model from http://ftp.ebi.ac.uk/pub/databases/alphafold/ and extracted the pLDDT score from these models for nine proteomes (UP000000437_7955_DANRE, UP000000803_7227_DROME, UP000002494_10116_RAT, UP000000589_10090_MOUSE UP000002195_44689_DICDI, UP000005640_9606_HUMAN, UP000000625_83333_ECOLI, UP000002311_559292_YEAST, and UP000006548_3702_ARATH). For each of these nine proteomes all PDB models were used, with the exception of the human proteome where only the first (F1) models were used. In total 151,970 different sequences were used.

**AlphaFold2 pLDDT predictor training**

The AlphaFold2 (AF2) pLDDT predictor (alphaPredict) was trained using PARROT (version 1.5.0), the same general-purpose deep learning framework used to create the BRNN behind the metapredict disorder predictions. The parameters used for PARROT training were as follows: residues datatype, 1 class (for regression), a learning-rate of 0.001, 2 hidden layers, 20 hidden vectors, a batch size of 32, and 200 training epochs. alphaPredict is being actively re-trained with larger datasets and more epochs to improve accuracy. To check the details of the most recent version check the metapredict documentation.

**AlphaFold2 pLDDT predictor accuracy**

At the time of writing, our AF2 pLDDT confidence score predictor obtained an $r^2$ value for the actual versus predicted scores in the test set was 0.7146 and the average error per residue was approximately 9.39%.

**AlphaFold2 pLDDT predictor implementation**

For improved modularity, the AlphaFold2 predictor is currently implemented as a separate Python package (alphaPredict) which is encoded as a silent dependency to metapredict (https://github.com/ryanemenecker/alphaPredict). alphaPredict can also be downloaded and used independently from metapredict in its own right. alphaPredict is written in Python 3.7+ and uses PyTorch, with the initial network trained using PARROT (Griffith & Holehouse, 2021; Paszke et al., 2019).

**AlphaFold2 pLDDT predictor improvements**

The prior information regarding the network (network V2) used for the AF2 pLDDT confidence score predictions in metapredict is up to date at the time of this writing. However, additional networks (a V3 and V4) are currently in progress. However, due to the increased number of proteomes used for training these networks and the slightly altered parameters, they currently have weeks of additional training remaining before completion. For the most up to date information on the AF2 pLDDT prediction in metapredict, please see https://github.com/ryanemenecker/alphaPredict.

### Supplemental Figures

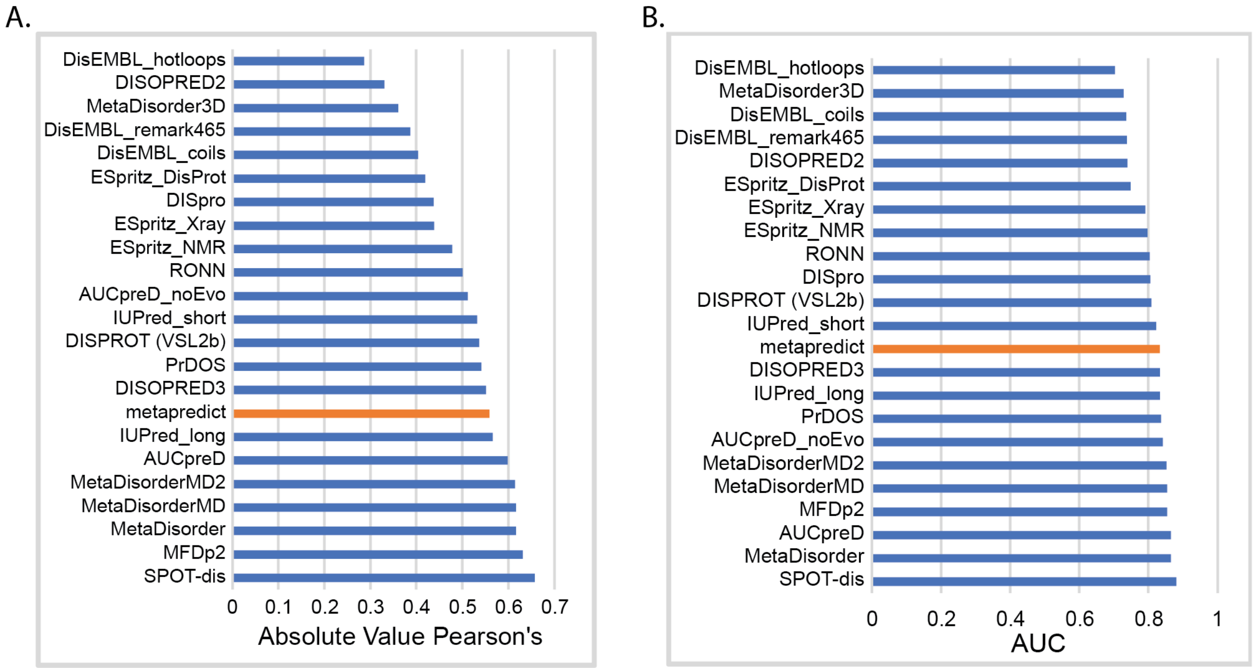


**Supplemental Figure 1. Evaluation of metapredict using CheZOD scores. (A)** The absolute value of the Pearson's correlation coefficient calculated by comparing the correlation between each predictor's score per residue and the CheZOD score. **(B)** The area under the receiver operating characteristic curve (AUC) (generated by comparing disorder scores of various predictors to disorder predictions from CheZOD scores. Values for all predictors in (A) and (B) other than metapredict (orange bar) were obtained from (Nielsen & Mulder, 2019).

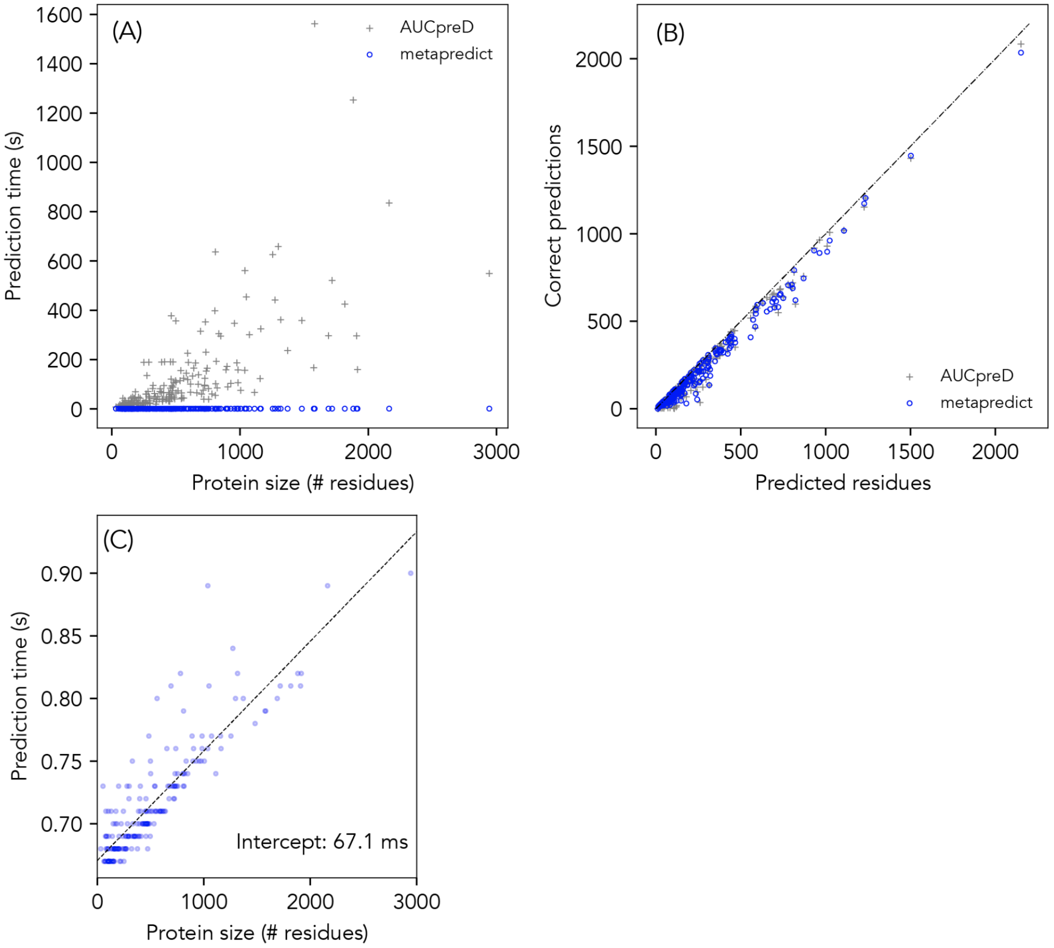


**Supplemental Figure 2. Comparison of execution time and predict performance for AUCPreD vs. metapredict (A)** Run time of 200 different proteins from the DISPROT dataset spanning a variety of different sizes as assessed by AUCPreD (grey crosses) vs. metapredict (blue circles). Both methods show a clear correlation between sequence length and execution time (AUCPreD Pearson's correlation coefficient of 0.71, metapredict Pearson's correlation coefficient of 0.88), yet the magnitude of the execution time for metapredict makes it look effectively flat. **(B)** Comparison of accuracy between AUCPreD and metapredict for the same sequences. The two methods are effectively comparable (see also **Fig. 3A**). **(C)** Zoomed-in comparison of execution time vs. protein size for metapredict. Note that the intercept here is 670 ms, which reflects the time needed to read-in and load the trained network, while the actual per-sequence execution time is under 100 ms for even a 1000 residue sequence (see also **Supplemental Fig. 8** where execution times are calculated after the initial network file has been read in and parsed by metapredict).

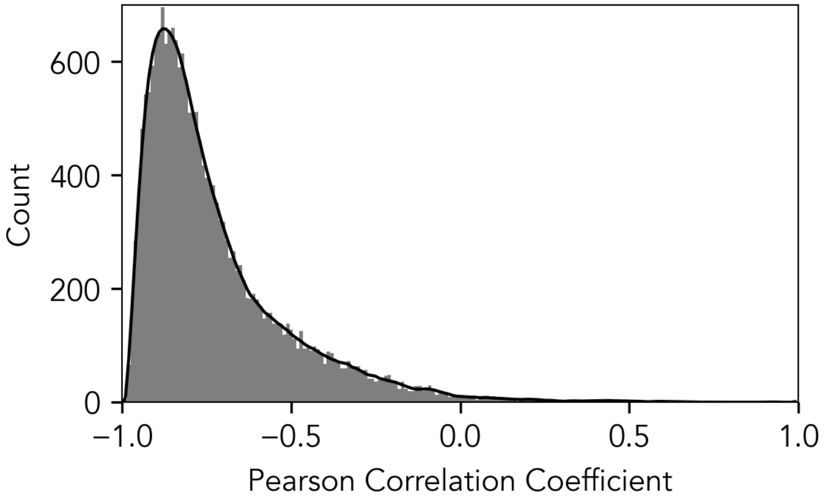


**Supplemental Figure 3. Disorder and predicted structure confidence are highly correlated but independent measures of (lack of) protein structure.**

We analyzed every sequence in the human proteome, computed per-residue disorder predictions and predicted structure confidence scores (predicted pLDDT), and correlated the scores using the Pearson correlation coefficient. The histogram above represents the overall distribution of those correlation coefficients calculated for 20,394 protein sequences.

A.

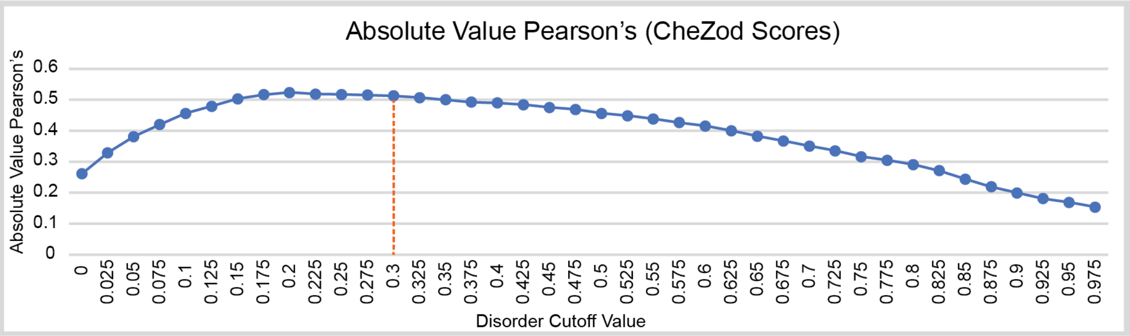


B.

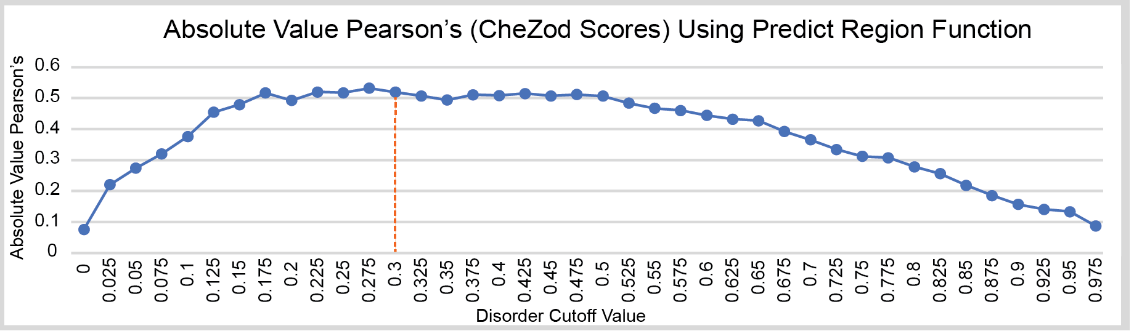


**Supplemental Figure 4. Assessing the impact of disorder cutoff values on binary order and disorder classification of CheZOD data. (A)** Absolute value of Pearson’s Correlation Coefficient for binary predictions of disorder by metapredict compared to binary classifications of disorder from the CheZOD dataset. **(B)** Absolute value of Pearson’s Correlation Coefficient for binary predictions of disorder by metapredict where the binary predictions were obtained using the predict_disorder_domains() function. These predictions are compared to binary classifications of disorder from CheZOD dataset. For both (A) and (B), the orange line represents the cutoff value used for binary classifications of order and disorder for metapredict.

A.

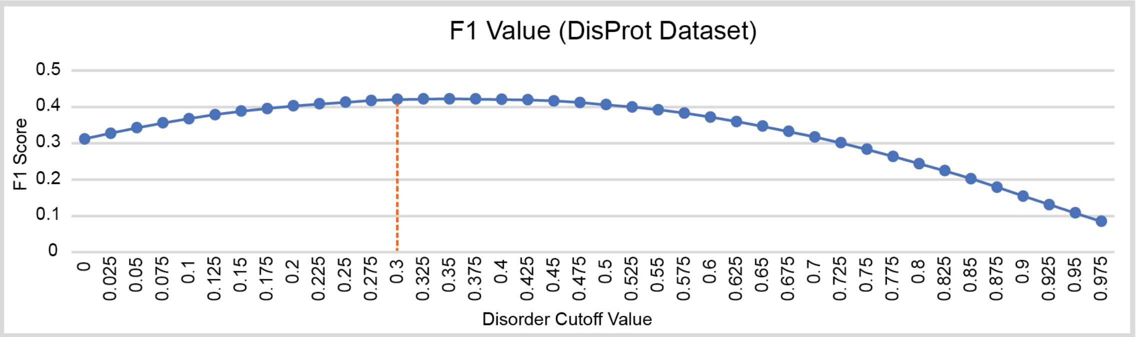


B.

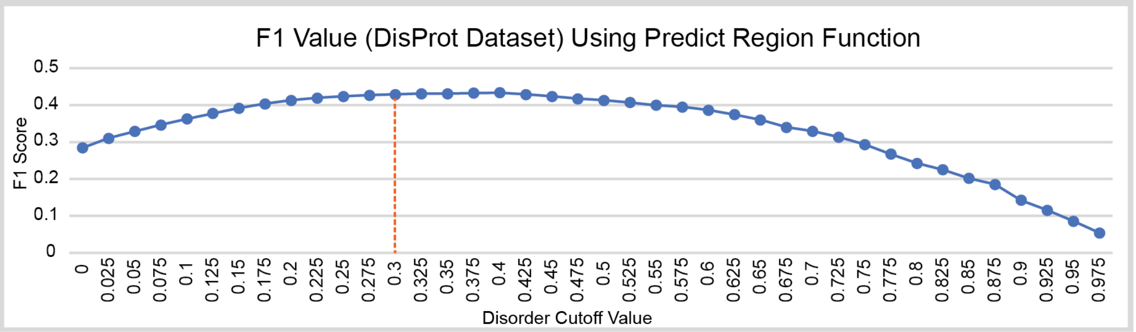


**Supplemental Figure 5. Assessing the impact of disorder cutoff values on binary order and disorder classification of the Disprot Dataset (CAID). (A)** F1-scores for binary predictions of disorder by metapredict compared to binary classifications of disorder from the Disprot dataset. **(B)** F1-scores for binary predictions of disorder by metapredict where the binary predictions were obtained using the predict_disorder_domains() function. These predictions were compared to binary classifications of disorder from the Disprot dataset. For both (A) and (B), the orange line represents the cutoff value used for binary classifications of order and disorder for metapredict.

A.

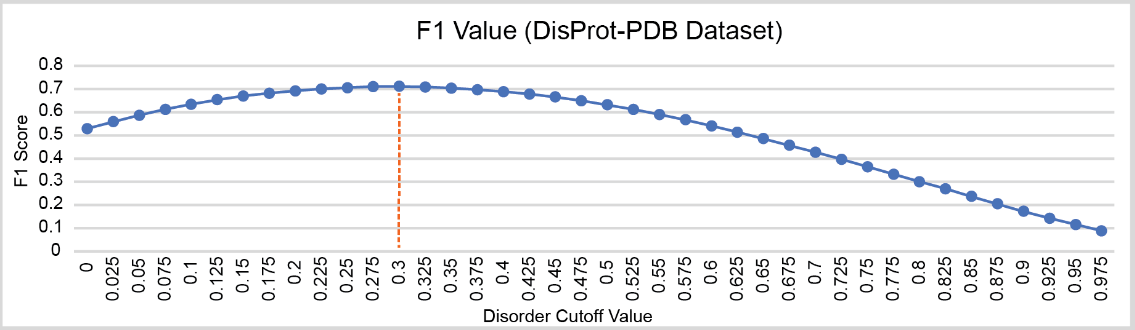


B.

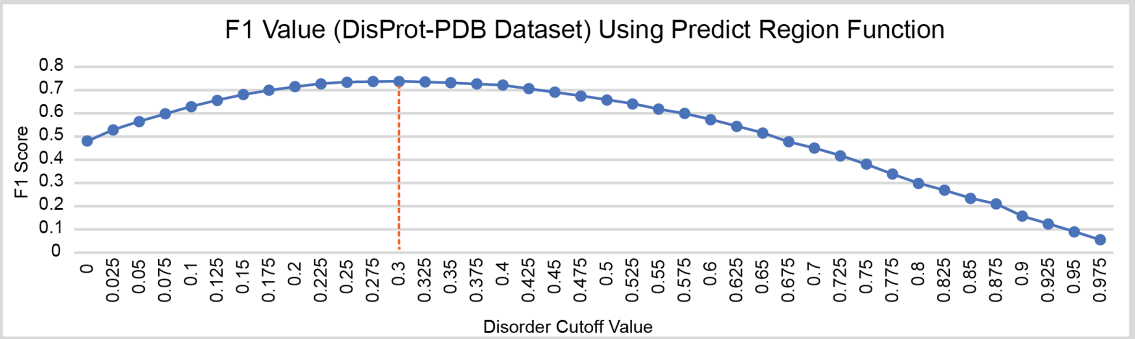


**Supplemental Figure 6. Assessing the impact of disorder cutoff values on binary order and disorder classification of the Disprot Dataset-PDB (CAID). (A)** F1-scores for binary predictions of disorder by metapredict compared to binary classifications of disorder from the Disprot-PDB dataset. **(B)** F1-scores for binary predictions of disorder by metapredict where the binary predictions were obtained using the predict_disorder_domains() function. These predictions were compared to binary classifications of disorder from the Disprot-PDB dataset. For both (A) and (B), the orange line represents the cutoff value used for binary classifications of order and disorder for metapredict.

A.

B.

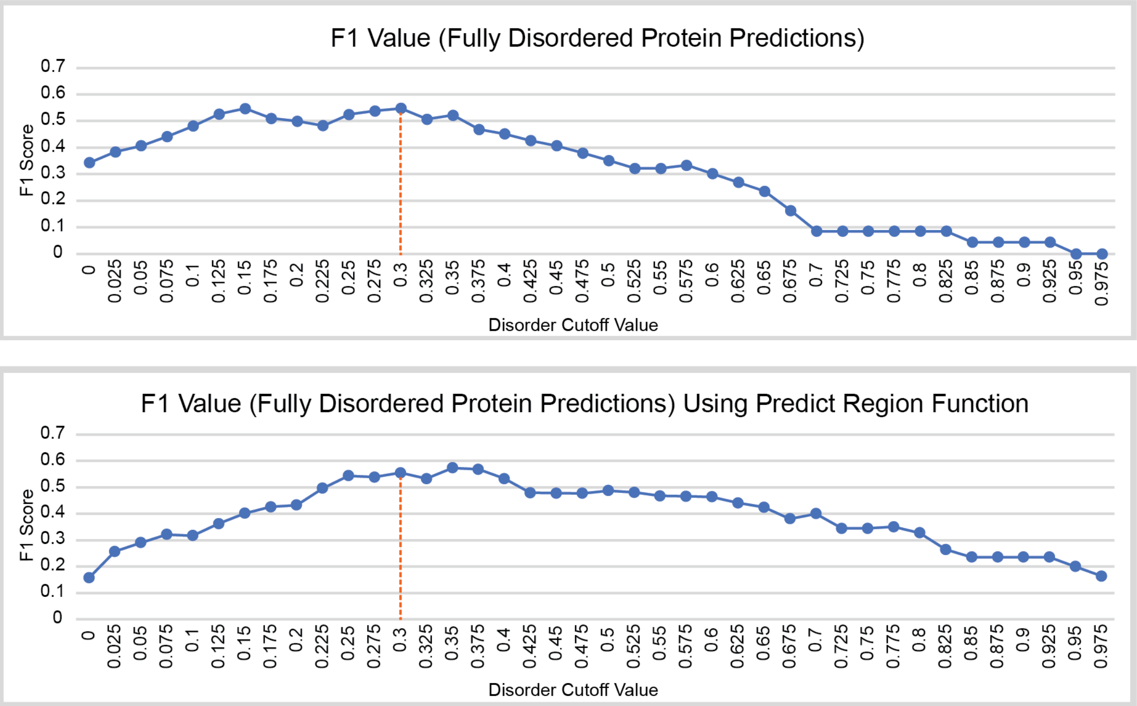


**Supplemental Figure 7. Assessing the impact of disorder cutoff values on metapredict identifying fully disordered proteins from the Disprot Dataset (CAID). (A)** F1-scores for predicted fully disordered proteins by metapredict compared to the known number of fully disordered proteins in the Disprot dataset. **(B)** F1-scores for predicted fully disordered proteins by metapredict where the binary predictions used to classify a protein as fully disordered were obtained using the predict_disorder_domains() function. These predictions were compared to the known number of fully disordered proteins in the Disprot dataset. For both (A) and (B), fully disordered proteins were counted if the predictor classified at least 95% of residues within a protein as disordered. For both (A) and (B), the orange line represents the cutoff value used for binary classifications of order and disorder for metapredict.

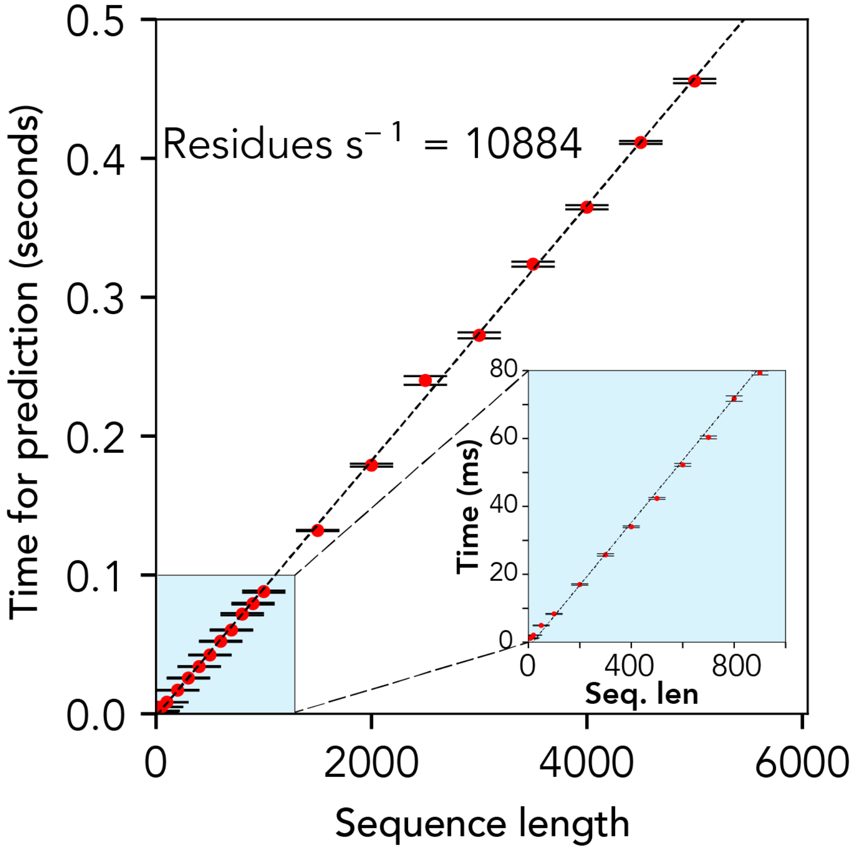


**Supplemental Figure 8. Metapredict performance as a function of sequence length in number of residues.** Assessment of length-dependence of metapredict performance reveals a linear scaling of prediction time with sequence length. Sequences here are randomly generated fixed-length sequences. Error bars are standard error of the mean calculated over thirty independent runs for random sequences of the specified length. Code for this analysis is provided in the Supporting Data GitHub repository.

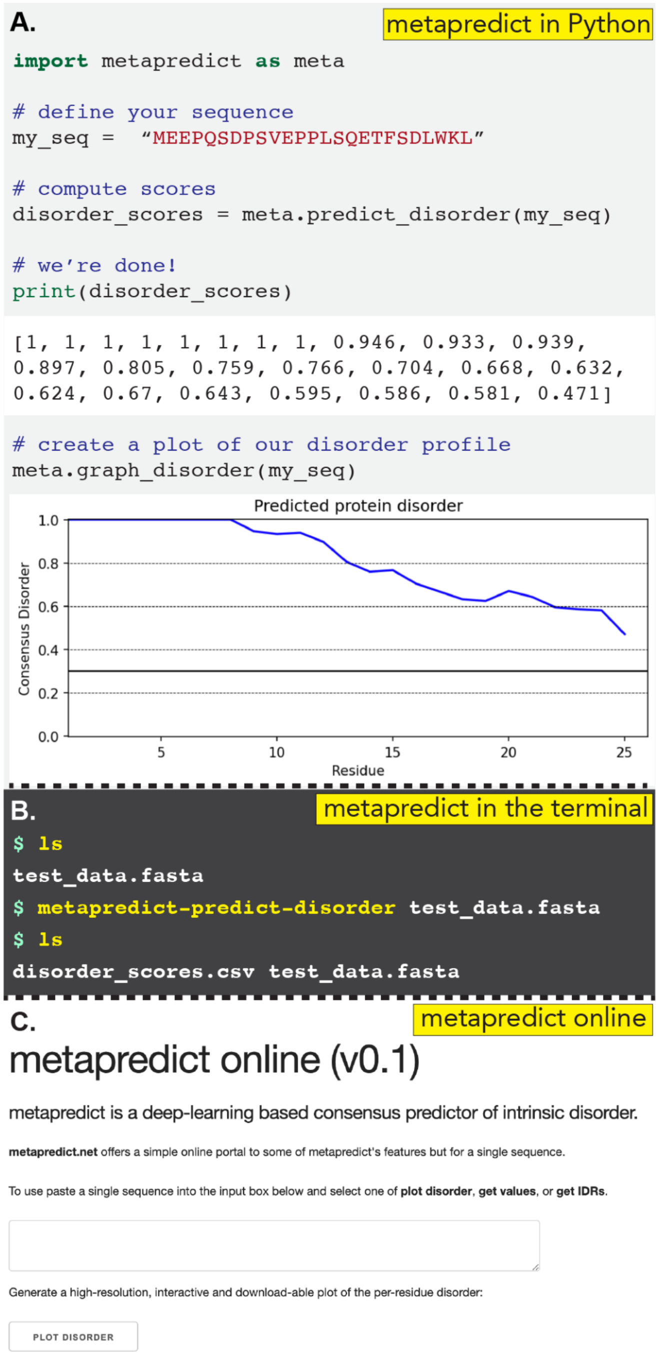


**Supplemental Figure 9. Metapredict offers three distinct modes of use. (A)** Metapredict can be used as a Python library, with simple and intuitive integration into existing Python code or for exploration in a Jupyter notebook. **(B)** Metapredict can be used as a command-line tool to interact directly with FASTA files. The file generated by the command metapredict-predict-disorder ("disorder_scores.csv") is a simple comma separated value (CSV) file with per-residue disorder values provided for each sequence in the FASTA file. **(C)** Finally, metapredict is offered as a simple web server (https://metapredict.net), which can generate high-quality downloadable figures or allow per-residue disorder scores to be obtained as a CSV data file.

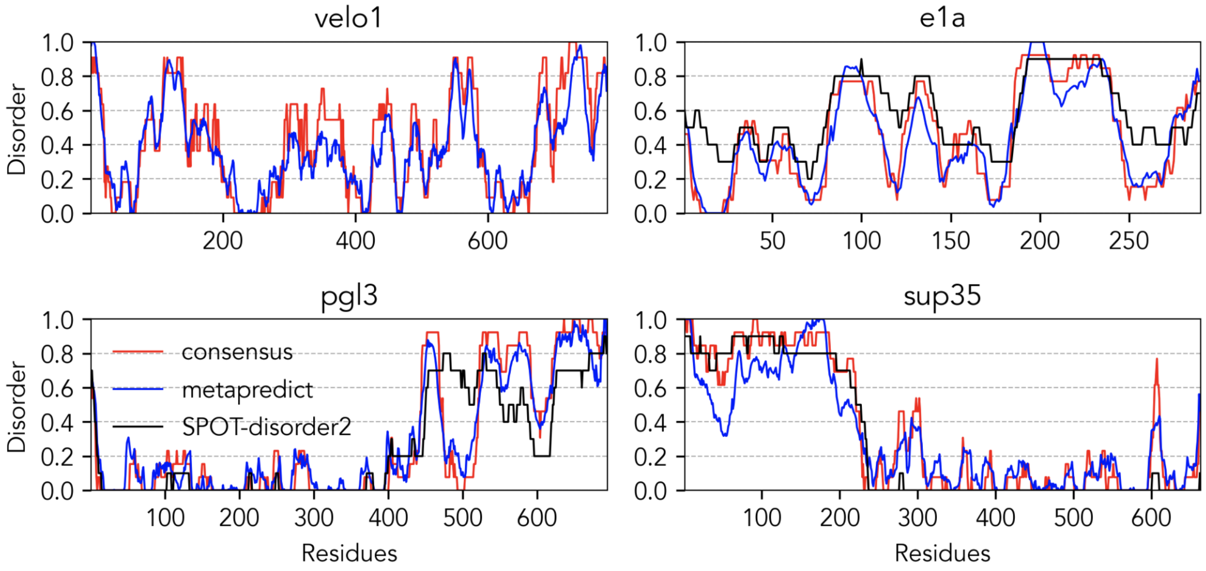


**Supplemental Figure 10. Metapredict accurately recapitulates precomputed consensus disorder scores.** Precomputed consensus disorder scores from the MobiDB database (red) and predicted disorder scores obtained SPOT-disorder2 (black) are compared to predicted consensus disorder scores calculated by metapredict for Velo1 from *Xenopus laevis* (UniProt Q7T226), PGL-3 from *Caenorhabditis elegans* (UniProt G5EBV6), Early E1A protein from Human adenovirus C serotype 5 (UniProt P03255), and Sup35 from *Schizosaccharomyces pombe* (UniProt O74718). None of these proteins were part of the training, test, or validation set for metapredicts. Note that Velo1 exceeds the length that SPOT-disorder2 can be used on.

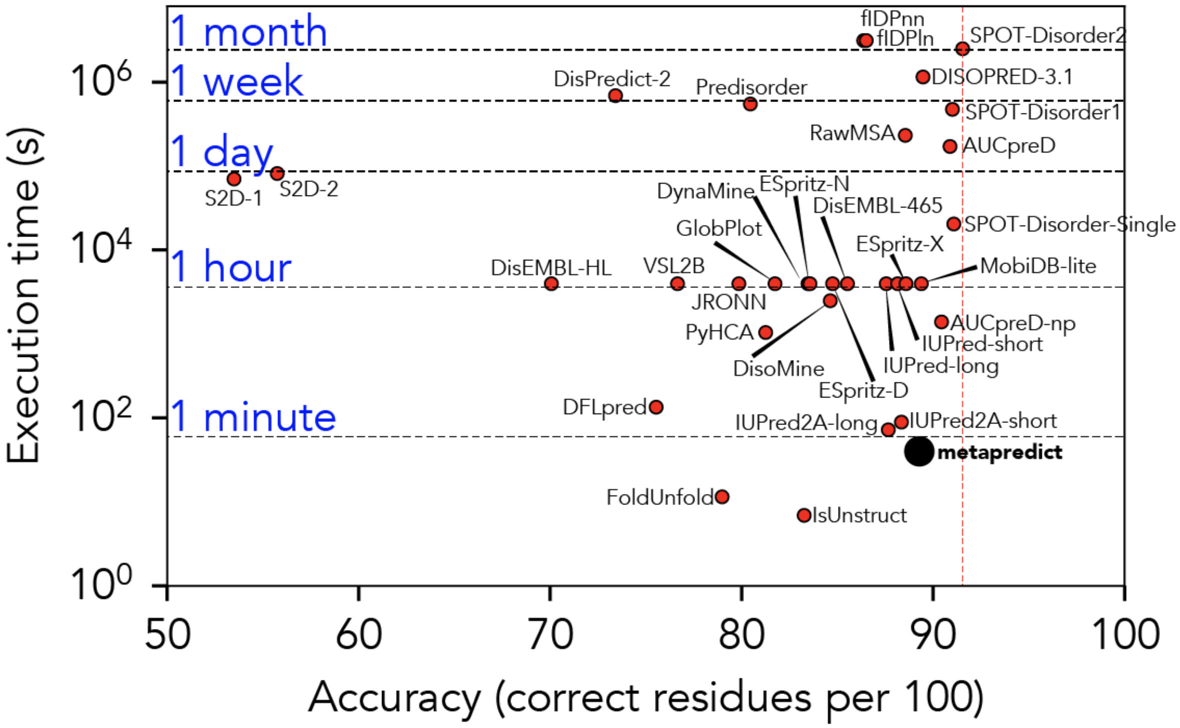


**Supplemental Figure 11.** Reproduction of Figure 3B with various predictors explicitly labelled.
